## Supplemental information for "Dynamic large-scale connectivity of intrinsic cortical oscillations supports adaptive listening in challenging conditions"

23562 Lübeck

**Dichotic listening task**

Participants listened to two competing, dichotically presented five-word sentences (Figure 1, main text). They were probed on the sentence-final noun in one of the two sentences. All sentences had the same structure and had an average length of 2512 ms (range: 2183–2963 ms). Sentences were spoken by the same female talker. Root mean (mean) square intensity (– 26 dB Full Scale, FS) was equalized across all individual sentences and they were masked by continuous speech-shaped noise at a signal-to noise-ratio of 0 dB. Noise onset was presented with a 50 ms linear onset ramp and preceded sentence onset by 200 ms. Each sentence pair was temporally aligned by the onset of the two task-related sentence-final nouns. All participants listened to the same 240 sentence pairs but in subject-specific randomized order. In addition, across participants we balanced the assignment of sentences to the right and left ear, respectively. Critically, two visual cues preceded auditory presentation. First, a spatial-attention cue either indicated the to-be-probed ear, thus invoking selective attention, or did not provide any information about the to-be-probed ear, thus invoking divided attention. Second, a semantic cue specified a general or a specific semantic category for the final word of both sentences, thus allowing to utilize a semantic prediction. Cue levels were fully crossed in a 2×2 design and presentation of cue combinations varied on a trial-by-trial level. Each trial started with the presentation of a fixation cross in the middle of the screen (jittered duration: mean 1.5 s, range 0.5–3.5 s; Figure 1, main text). Next, a blank screen was shown for 500 ms followed by the presentation of the spatial cue in the form of a circle segmented equally into two lateral halves. In selective-attention trials, one half was black, indicating the to-be-attended side, while the other half was white, indicating the to-be-ignored side. In divided-attention trials, both halves appeared in grey. After a blank screen of 500 ms duration, the semantic cue was presented in the form of a single word that specified the semantic category of both final words. The semantic category could either be given at a general (natural vs. man-made) or specific level (e.g. instruments, fruits, furniture) and thus provided different degrees of semantic predictability. Each cue was presented for 1000 ms. After a 500 ms blank-screen period, the two sentences were presented dichotically along with a fixation cross displayed in the middle of the screen. Finally, after a jittered retention period, a visual response array appeared on the left or right side of the screen, presenting four word-choices. The location of the response array indicated which ear (left or right) was probed. Participants were instructed to select the final word presented on the to-be-attended side using the touch screen. Among the four alternatives were the two actually presented nouns as well as two distractor nouns from the same cued semantic category. Note that because the semantic cue applied to all four alternative verbs, it could not be used to post-hoc infer the to-be-attended final word. Stimulus presentation was controlled by PsychoPy. The visual scene was displayed using a 24" touch screen (ViewSonic TD2420) positioned within an arm's length. Auditory stimulation was delivered using in-ear headphones (EARTONE 3A) at sampling rate of 44.1 kHz. Following instructions, participants performed a few practice trials to familiarize themselves with the listening task. To account for differences in hearing acuity within our group of participants, individual hearing thresholds for a 500-ms fragment of the dichotic stimuli were measured using the method of limits. All stimuli were presented 50 dB above the individual sensation level. During the experiment, each participant completed 60 trials per cue-cue condition, resulting in 240 trials in total. The cue conditions were equally distributed across six blocks of 40 trials each (~ 10 min) and were presented in random order. Participants took short breaks between blocks.

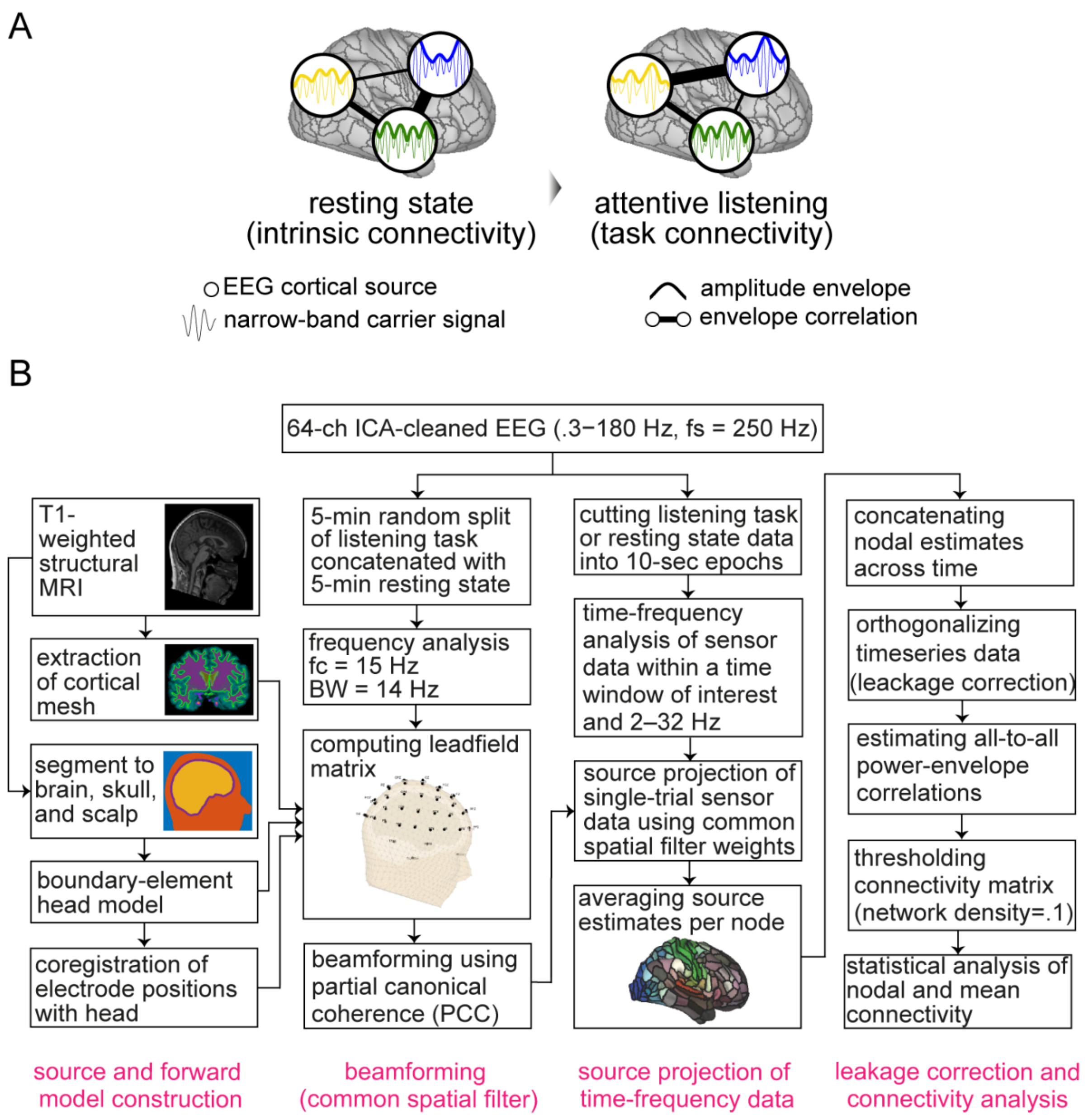


**Figure S1** Hypothetical and methodological framework **(A)** Toy graphs illustrate connectivity between three exemplary EEG cortical sources as measured by power-envelope correlation. During attentive listening intrinsic neural oscillations regulate their putative baseline network (resting state) depending on the current task state to support listening behavior. These dynamics could manifest as change in connectivity strength (here depicted as edge thickness) or network segregation [5]. **(B)** EEG source analysis pipeline. Source reconstruction of EEG oscillatory responses and estimation of connectivity per individual (N = 154) were implemented in four steps: (1) construction of source geometry using MRI T1-derived cortical mesh and head boundary-element model (2) estimation of lead-field matrix and a spatial filter common across rest, task, and a broad frequency band (center frequency (fc) = 15 Hz; band-width (BW) = 14 Hz) (3) source projection of time-frequency sensor data and averaging the source estimates per cortical parcel or node (4) pair-wise orthogonalization of complex-value time-frequency source estimates across all nodes to diminish spurious correlations due to volume conduction [25]. These steps at the end gave individual cortical connectivity maps per time-window of interest and frequency band. These analyses were performed using Fieldtrip toolbox for Matlab and Human Connectome Project (HCP) functional parcellation template [46].

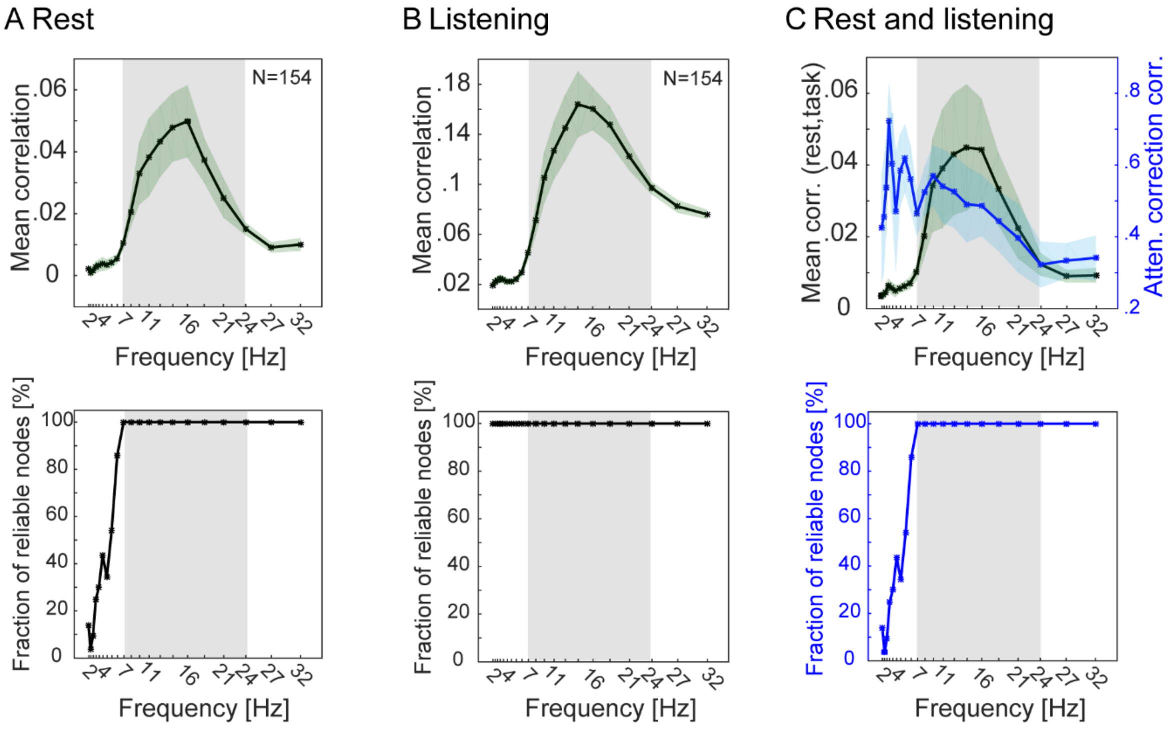


**Figure S2** Between-subject reliability of power-envelope correlation.

A serious but often neglected potential confounding factor in assessments of neuronal interactions is signal-to-noise ratio (SNR) [48, 45]. This is an important issue in the present study as we compare connectivity of frequency-specific neural oscillations between resting state and listening task, each of which is potentially measured at different levels of SNR. Thus, we first investigated at which frequencies connectivity can be reliably measured under both rest and task conditions. As a measure of reliability, we used between-subject correlation of rest and task nodal connectivity values, i.e., columns of raw connectivity matrices. The reliability analysis was done in two steps. **(A/B)** First, per cortical node, correlations between nodal connectivity values were calculated across all pairs of N = 154 participants, separately for rest or task condition. This procedure per node gives one symmetric N×N correlation matrix for each rest or task condition. The upper-diagonal average of this matrix is mean between-subject correlation in nodal connectivity for each rest or task condition (within-condition reliability). The plots in (A) and (B) illustrate mean ± SEM of these between-subject correlations averaged across all 360 cortical nodes per frequency. **(C)** Next, the same analysis was done across rest and task. In this case the result per node is one asymmetric N×N correlation matrix. The off-diagonal average of this matrix is mean between-subject correlation in nodal connectivity across rest and task (between-condition reliability). The black line graph in (C) illustrates mean ± SEM of these between-subject correlations averaged across all 360 cortical nodes per frequency. These correlations are underestimated in the presence of noise. Thus, they were submitted to Spearman's correction for attenuation to account for differences in SNR between rest and task (blue line graph). Estimation of connectivity at a given frequency and node was considered reliable if the attenuation-corrected correlation was consistently positive across all participants. Plots in second row illustrate percentage of cortical nodes showing significantly positive between-subject correlation ($p<0.05$, uncorrected). As illustrated by both blue line graphs, between-subject correlations were consistently positive across all nodes within the frequency range 7-32 Hz ($0.1 < r < 0.4$). These results illustrate that power-envelope correlations between EEG oscillatory sources can be reliably measured within alpha and low-beta frequency range (grey frequency intervals).

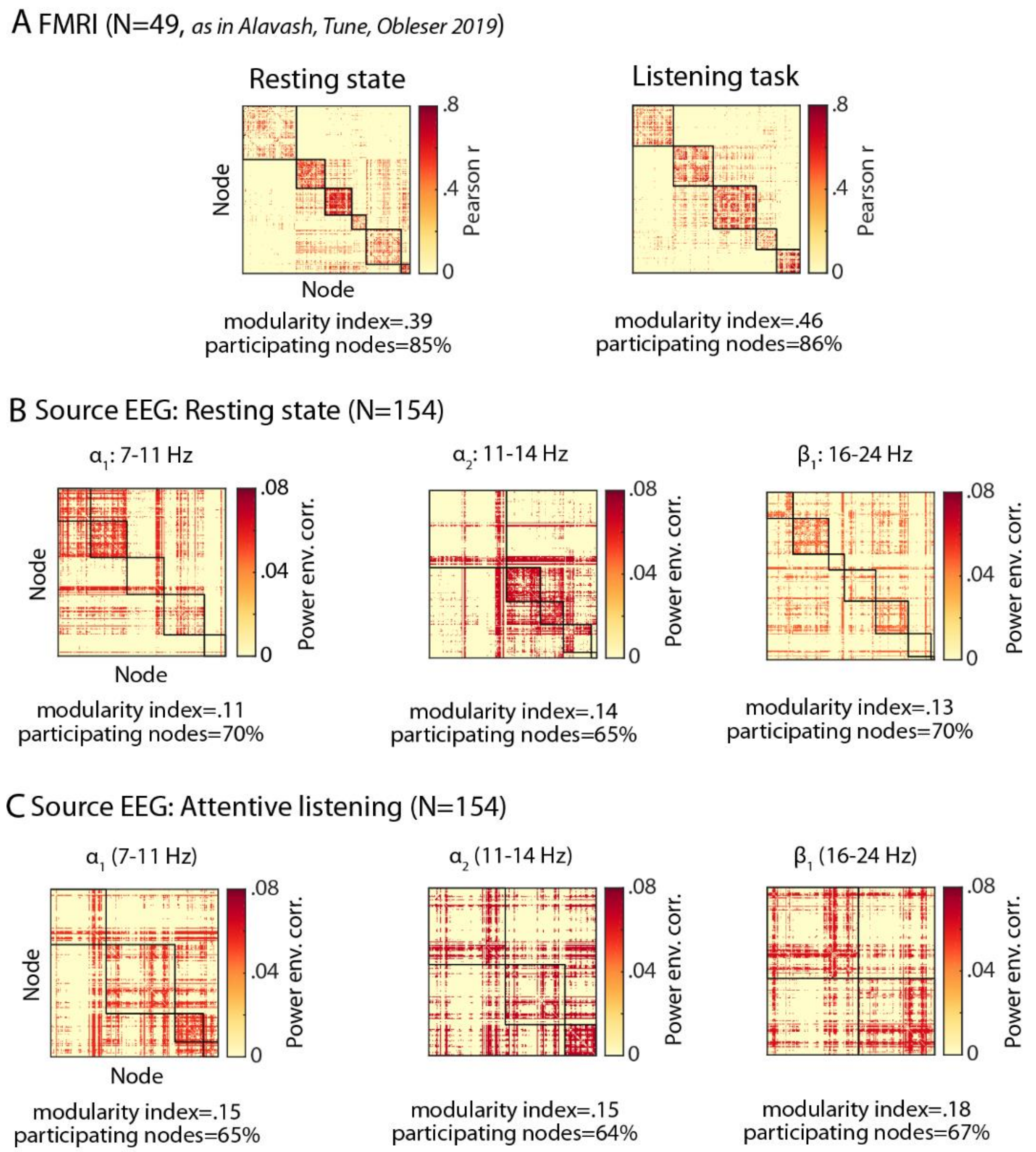


**Figure S3** Network modularity of α/β oscillations under rest and attentive listening. In each group-average connectivity matrix, diagonal squares represent functional modules as detected by Newman algorithm. Off-diagonal correlations represent between-module connections. Quantitatively, the modular structure of a network can be estimated by optimizing the so-called modularity index (Blondel et al., 2008; Rubinov and Sporns, 2010): The more modular the network, the closer the index to one. In A, functional modules correspond to canonical resting state networks (left) and their makeup during the listening task (right, see Alavash et al., 2019). In contrast, EEG source power-envelope correlations within alpha and low-beta bands did not exhibit a modular organization. Connectivity matrices are averaged across individuals and thresholded at 10% of network density. Modularity index ($Q$) quantifies the degree to which a network is clustered into densely intra-connected groups of nodes which are sparsely inter-connected and is estimated based on Newman optimization algorithm [44, 45] (the more modular the network, the closer the $Q$ to 1). Percentage of participating nodes is calculated as the number of nodes having at least one connection in the network divided by the total number of nodes defined according to the cortical parcellation template [46].

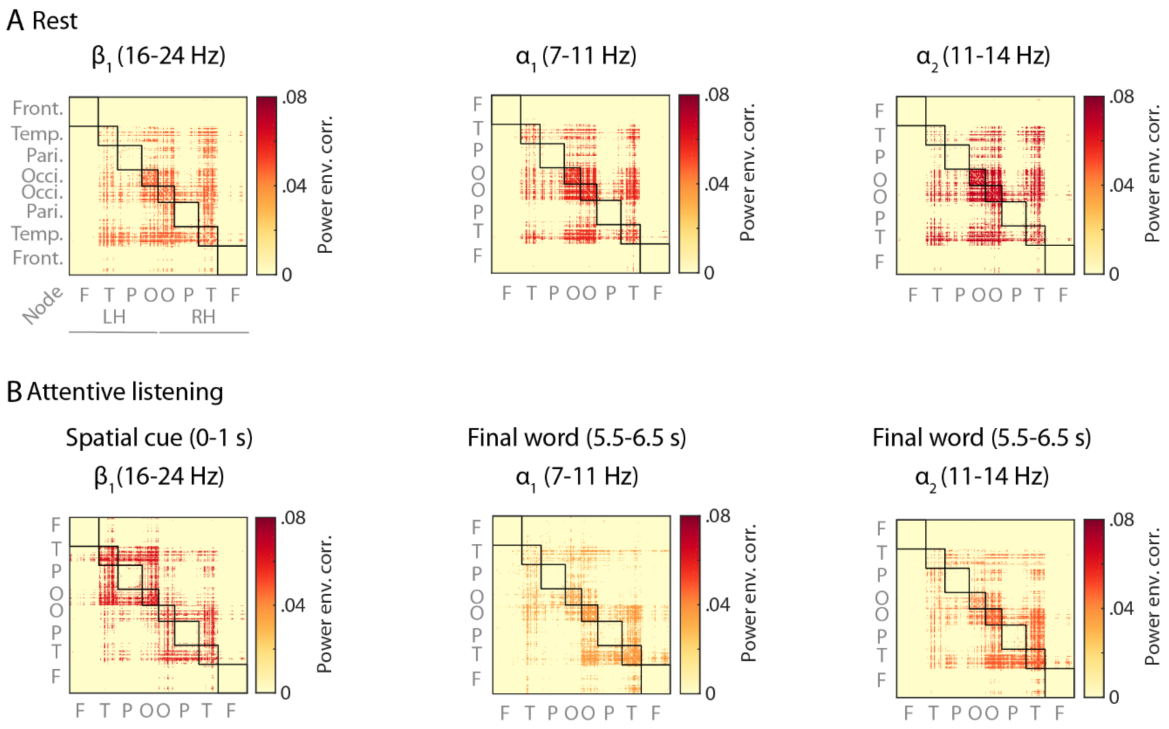


**Figure S4** Frequency-specific connectivity matrices **(A)** For each frequency band, power-envelope correlations between EEG oscillatory sources were estimated using 4-min eyes-open resting state data **(B)** Whole-brain connectivity during task time intervals most critical to listening behavior. Power-envelope correlations were estimated by concatenating 1-s windowed signals across all 240 trials (4-min data). Note the stronger beta-band connectivity during processing of spatial cue and weaker alpha-band connectivity during final word presentation in (B) relative to resting state (A). Connectivity maps were averaged across N = 154 individuals and thresholded at 10% of network density. Nodes correspond to cortical parcels as in [46] and are grouped according to their cortical lobes. *LH*: left hemisphere; *RH*: right hemisphere.

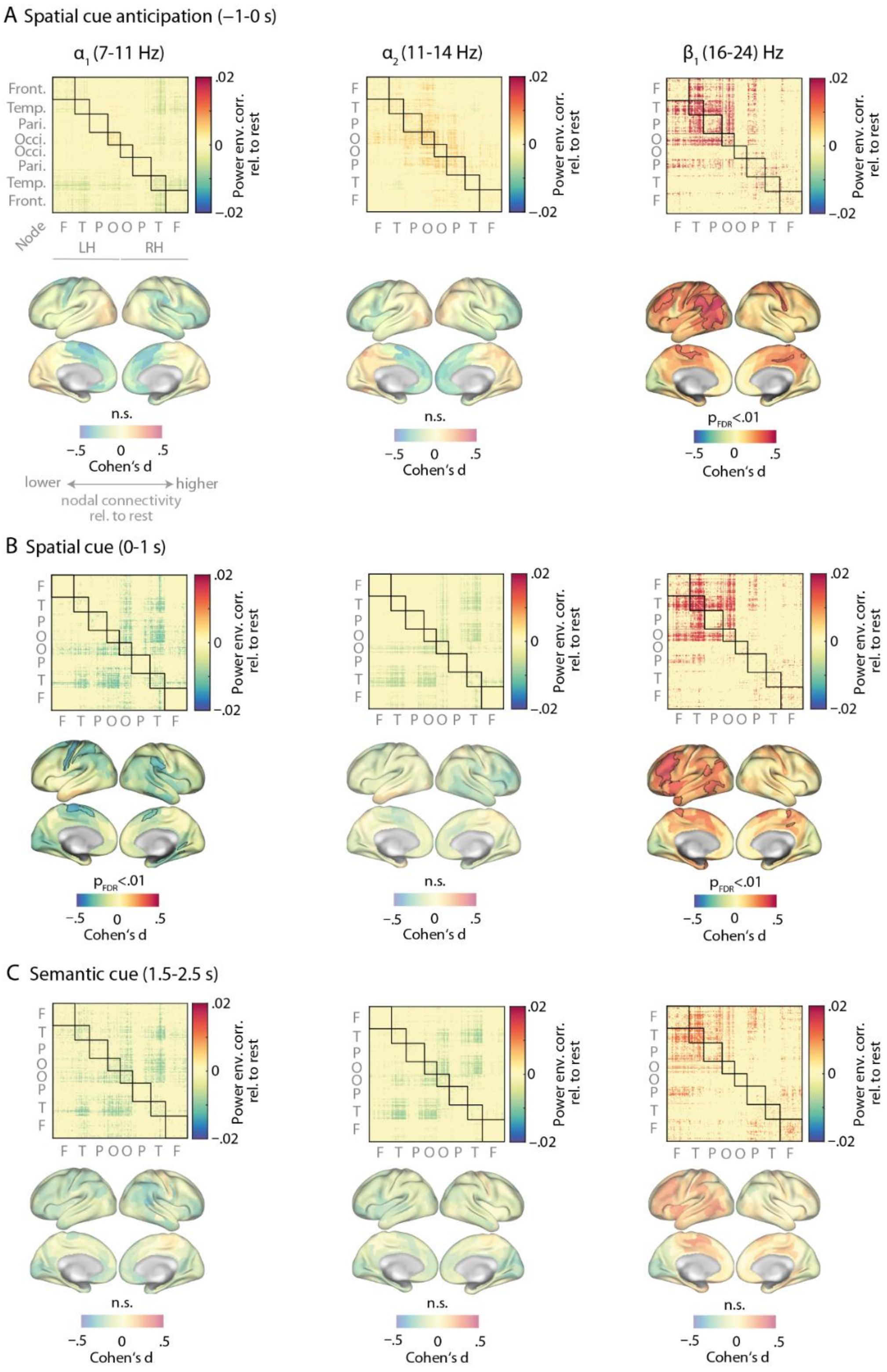


**Figure S5** Cortical connectivity dynamics of alpha and low-beta oscillations during anticipation and cueing. For each time interval (A-C) and frequency band, power-envelope correlations between EEG oscillatory sources were estimated by concatenating 1-s windowed signals across all 240 trials (4-min data) and compared with 4-min resting state connectivity at the same frequency band (i.e., task minus rest). In anticipation of and during the spatial cue presentation $\boldsymbol{\beta_1}$ connectivity was significantly increased mainly across frontoparietal regions (A/B, third column; paired-sample permutation tests, significant nodes are outlined in black). Alpha-band connectivity was not significantly different than rest. Connectivity difference maps were averaged across N = 154 individuals and thresholded at 10% of network density. Nodes correspond to cortical parcels as in [46] and are grouped according to their cortical lobes. *LH*: left hemisphere; *RH*: right hemisphere.

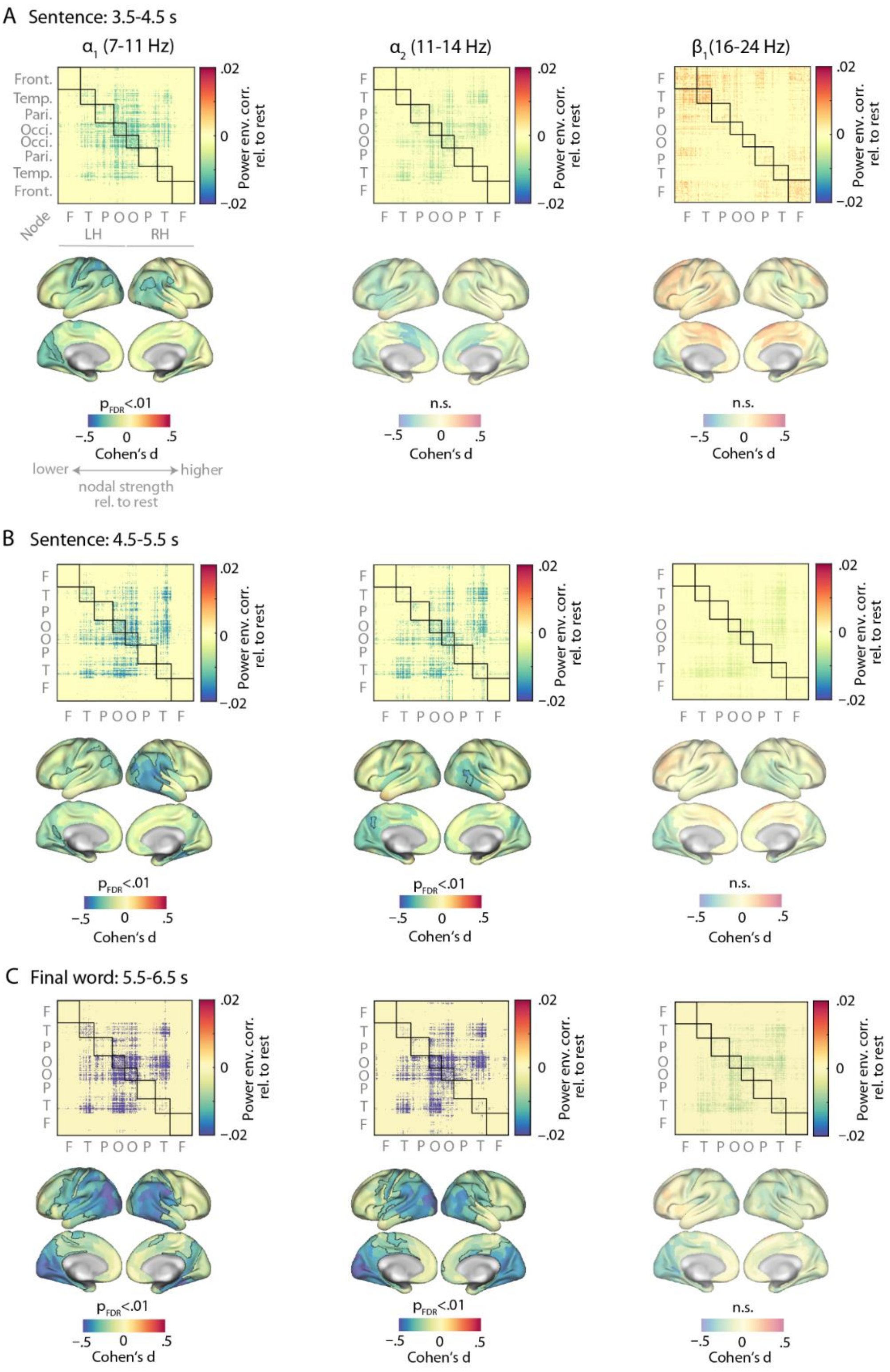


**Figure S6** Cortical connectivity dynamics of alpha and low-beta oscillations during sentence presentation. For each time interval of sentence presentation (A-C) and frequency band, power-envelope correlations between EEG oscillatory sources were estimated by concatenating 1-s windowed signals across all 240 trials (4-min data) and compared with 4-min resting state connectivity at the same frequency band (i.e., task minus rest). Towards the end of sentence and particularly during final word presentation alpha-band connectivity was significantly decreased across posterior cortical regions (A-C, first column; paired-sample permutation tests, significant nodes are outlined in black). Beta-band connectivity was not significantly different than rest during sentence presentation. Connectivity difference maps were averaged across N = 154 individuals and thresholded at 10% of network density. Nodes correspond to cortical parcels as in [46] and are grouped according to their cortical lobes. *LH*: left hemisphere; *RH*: right hemisphere.

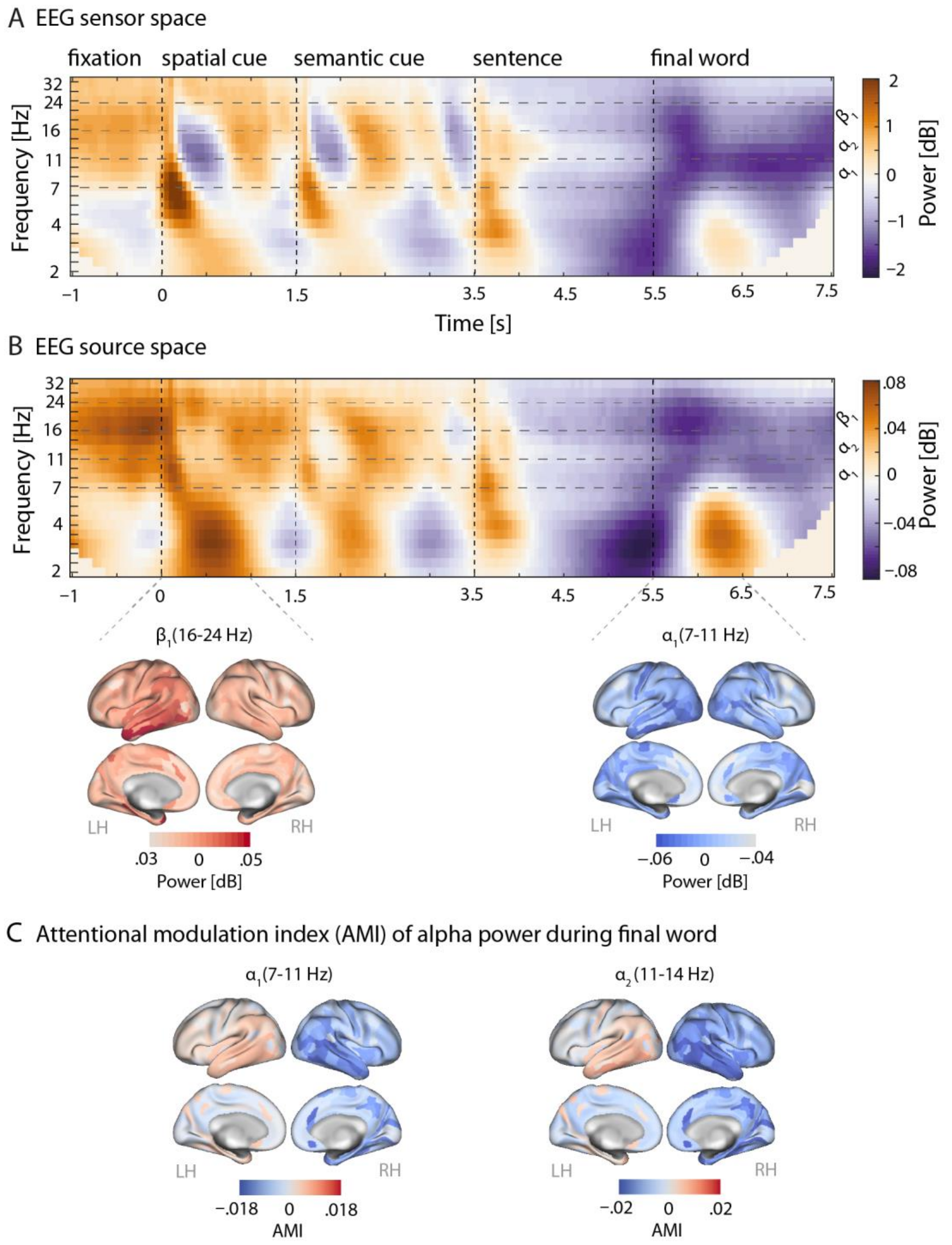


**Figure S7** Modulation of neural oscillatory activity during attentive listening. **(A)** Time-frequency representation in EEG sensor space. Power estimates were averaged across all sensors, trials, and N = 156 participants. **(B)** The same as in (A) but based on power of EEG source estimates and averaged across all cortical nodes. Power changes are calculated as dB change relative to the whole-trial baseline interval (–1-7.5 s). **(C)** Attentional modulation of alpha source power during selective attention to final word. The modulation index is calculated as AMI = ($\alpha$-power$_{attendL}$ – $\alpha$-power$_{attendR}$) / ($\alpha$-power$_{attendL}$ + $\alpha$-power$_{attendR}$). *LH*: left hemisphere. *RH*: right hemisphere.

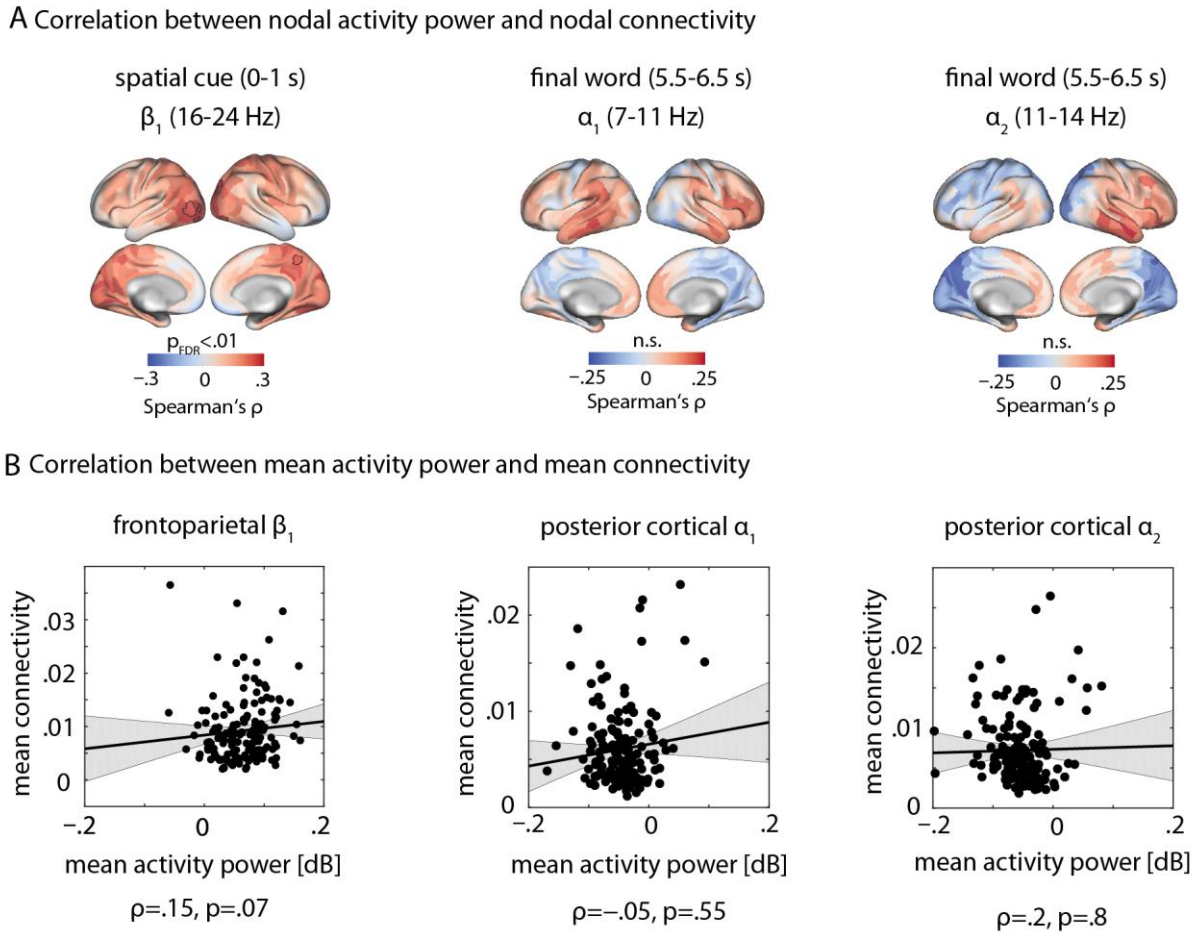


**Figure S8** Correlation between neural activity power and connectivity during time intervals of listening task. (A) For each frequency band and time interval, correlation between nodal connectivity and mean nodal power (dB) was tested across all N = 154 participants and per cortical node. (B) The same analysis as in (A) but using measures averaged across frontoparietal or posterior cortical nodes involved in beta-band hyperconnectivity or alpha-band hypoconnectivity, respectively.

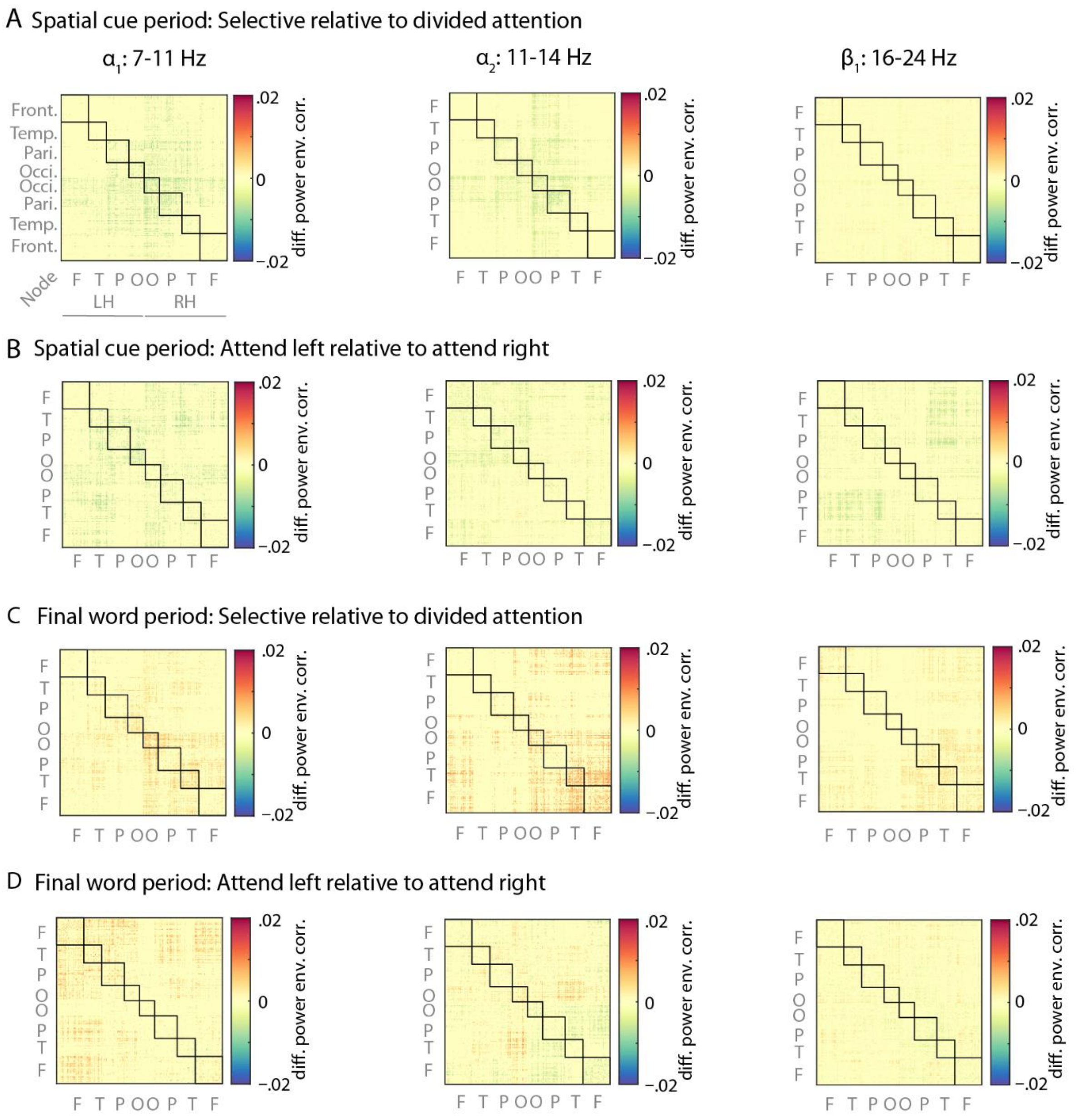


**Figure S9** EEG source connectivity dynamics were not influenced by spatial cue conditions. We investigated whether alpha- or beta-band connectivity showed a similar hemispheric lateralization in response to the spatial cue as in alpha power. For each spatial-cue (A/B) or final-word (C/D) time interval and frequency band, power-envelope correlations between EEG oscillatory sources were estimated by concatenating 1-s windowed signals across trials per spatial-cue condition: selective attention (i.e., attend left or right), divided attention, attend left, or attend right. The results were then compared by first calculating connectivity difference maps per participant (e.g., selective minus divided) and then averaging the difference maps across N = 154 individuals. If connectivity is confounded by activity level, these maps should reveal differences in power-envelope correlations between conditions. In contrast, power-envelope correlations were not influenced by attentional-cue conditions in neither of the trial intervals nor frequency bands. None of the connectivity contrasts revealed significant differences in nodal connectivity when tested across all cortical nodes using paired-sample permutation tests. Nodes correspond to cortical parcels as in [46] and are grouped according to their cortical lobes. *LH*: left hemisphere; *RH*: right hemisphere.

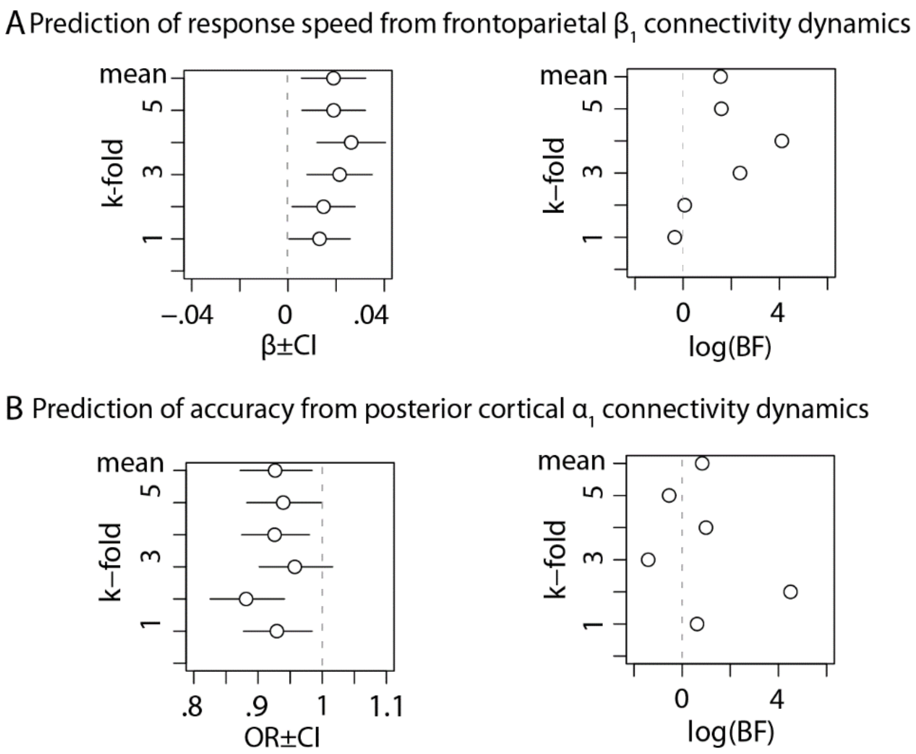


**Figure S10** Within-sample reproducibility and likelihood assessment of the brain-behavior interaction effects**.** The reliability and robustness of the brain-behavior interaction effects (main text, Fig. 5) were investigated by randomly splitting the data into k = 5 non-overlapping folds. The strength and uncertainty (i.e., confidence interval) of the model's parameter estimate was then examined when each fold of data was used. Since this analysis does not take the number of observations and model complexity into account, it could be that the model has a relatively high goodness-of-fit as a too complex model is fitted to the data. We thus examined the relative strength of evidence (rather than statistical significance) in support of the alternative hypothesis (i.e., model with the interaction term) as compared to the null hypothesis (i.e., model without the interaction term). This was done by calculating the BIC approximation of the Bayes factor, i.e., exp([BIC(H0)–BIC(H1)]/2). In this calculation BIC imposes penalty on each model log-likelihood based on both number of observations and parameters. Following Harrold Jeffery's scale for interpretation of BF, 0 < log-BF < 1 suggests weak evidence for H1, 1 < log-BF < 3 is positive, and 3 < log-BF < 5 is strong [106]. *β*: Slope parameter estimates from linear mixed-effects model. *OR*: Odds ratio parameter estimates from generalized linear mixed-effects models.

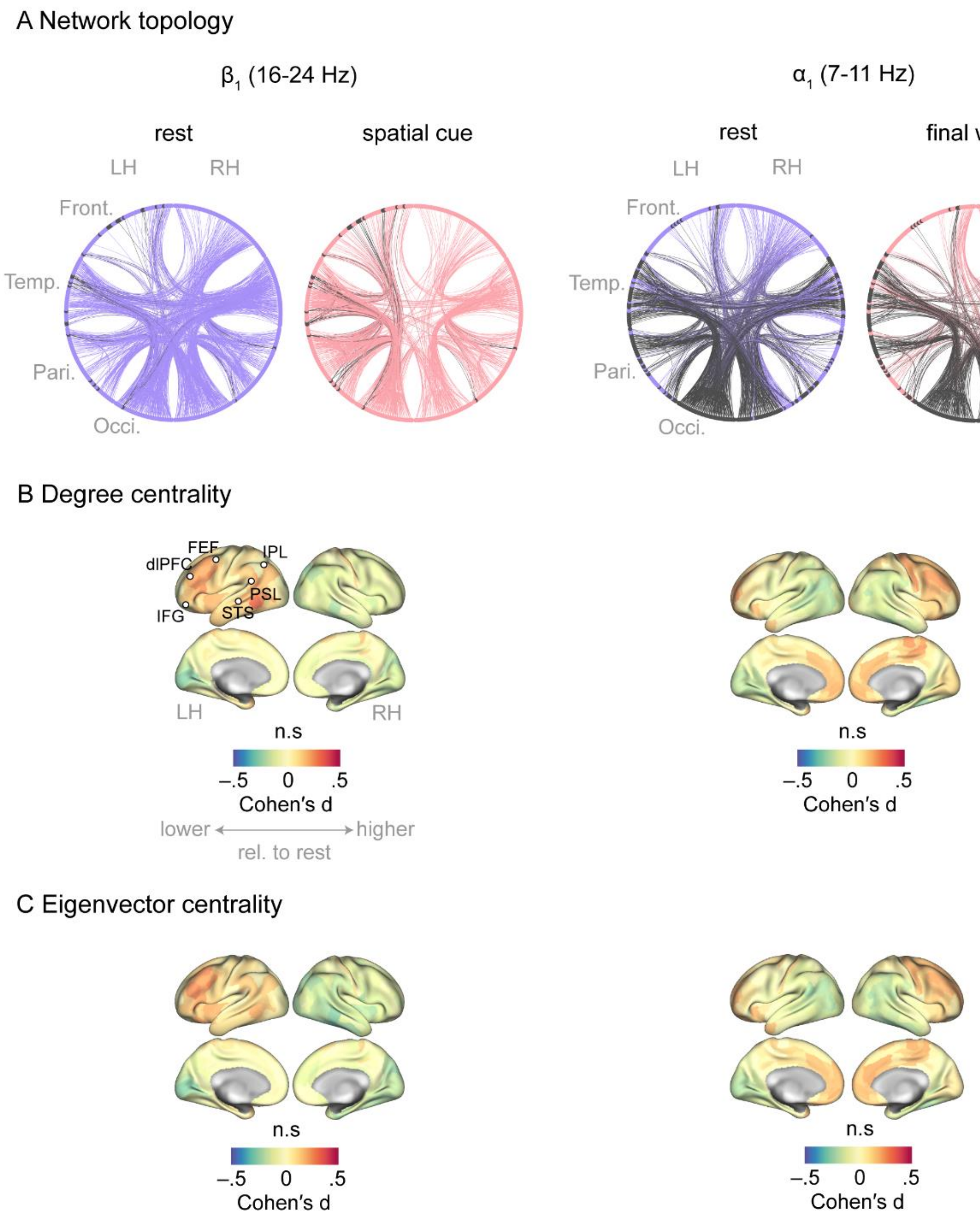


**S11 Fig.** Network topology and centrality of α/β oscillations under rest and attentive listening**. (A)** Network topology per frequency band and condition. Each connectogram corresponds to its connectivity matrix counterpart and is obtained by setting the connection weights to 1 for correlations higher than 95-th percentile, and to 0 otherwise (i.e., a sparse binary graph thresholded at 5% of density). Nodes correspond to cortical parcels as in [46] and are grouped according to their cortical lobes. The frontoparietal and posterior cortical nodes showing significant modulation of $\beta_1$ and $\alpha_1$ connectivity strength, respectively (main text, Fig. 5), are depicted in black. **(B)** Comparison of nodal degree centrality between each task interval and rest using permutation tests. Degree centrality is defined as each node's number of connections (also referred to as node's degree [44]). This measure quantifies relative importance of nodes: nodes having large number of connections, hence high degree centrality, can be considered as central coordinators in the functional network. In contrast to nodal connectivity strength, nodal degree was not significantly modulated when tested across all cortical nodes (brain surfaces; significance level: $p_{FDR}<0.01$). **(C)** Comparison of eigenvector centrality. This metric is more specific than degree centrality as it also takes the relative importance of a node's neighbors into account. Eigenvector centrality of node $i$ is equivalent to the $i$th element in the eigenvector corresponding to the largest eigenvalue of the adjacency matrix. Nodes having high eigenvector centrality are high-degree nodes whose neighbors also have relatively large number of connections. Eigenvector centrality was not significantly modulated when tested across all cortical nodes. *LH*: left hemisphere, *RH*: right hemisphere, *STS*: superior temporal sulcus, *PSL*: perisylvian language area, *IPL*: inferior parietal lobule, *FEF*: frontal eye field, *dlPFC*: dorsolateral prefrontal cortex, *IFG*: inferior frontal gyrus.

**Table S1** Summary table of the generalized linear mixed-effects model predicting individuals' listening accuracy. In this model brain regressors were based on the data during final word period and within $\boldsymbol{\alpha_1}$ band. Significant effects after FDR-correction for multiple comparisons across model terms are shown in blue. The main effects of and interactions between listening cues are visualized in Figure 2 (main text). The interaction between rest and task connectivity is visualized in Figure 5. *OR*: Odds ratio; $\boldsymbol{\sigma^2}$: within-group variance; $\boldsymbol{\tau_{00}}$: between-group variance; $\boldsymbol{\rho_{01}}$: random-slope-intercept-correlation.

| | Accuracy | | | | |
|---|---|---|---|---|---|
| Predictors | Odds Ratios | std. Error | CI | Statistic | $p_{FDR}$ |
| (Intercept) | 20.650 | 0.088 | 17.362 – 24.561 | 34.213 | <0.001 |
| AttnCueSelective | 3.438 | 0.118 | 2.725 – 4.337 | 10.421 | <0.001 |
| SemCueSpecific | 1.107 | 0.107 | 0.897 – 1.367 | 0.948 | 0.343 |
| ProbeRight | 1.249 | 0.068 | 1.093 – 1.426 | 3.275 | 0.001 |
| Age | 0.788 | 0.082 | 0.671 – 0.924 | -2.925 | 0.003 |
| PTA | 0.763 | 0.080 | 0.652 – 0.894 | -3.358 | 0.001 |
| Block | 1.284 | 0.021 | 1.233 – 1.337 | 12.036 | <0.001 |
| FinalWordAlpha1Conn | 0.986 | 0.035 | 0.921 – 1.056 | -0.395 | 0.693 |
| RestAlpha1Conn | 0.910 | 0.074 | 0.788 – 1.052 | -1.274 | 0.203 |
| FinalWordAlpha1Pow | 0.963 | 0.047 | 0.879 – 1.056 | -0.791 | 0.429 |
| AttnCueSelective : SemCueSpecific | 1.360 | 0.215 | 0.892 – 2.071 | 1.430 | 0.153 |
| AttnCueSelective * FinalWordAlpha1Conn | 1.004 | 0.055 | 0.902 – 1.119 | 0.081 | 0.935 |
| AttnCueSelective * RestAlpha1Conn | 0.999 | 0.066 | 0.879 – 1.137 | -0.008 | 0.993 |
| AttnCueSelective * FinalWordAlpha1Pow | 1.024 | 0.057 | 0.915 – 1.145 | 0.407 | 0.684 |
| SemCueSpecific * FinalWordAlpha1Conn | 1.041 | 0.043 | 0.956 – 1.132 | 0.924 | 0.356 |
| SemCueSpecific * RestAlpha1Conn | 0.983 | 0.044 | 0.902 – 1.072 | -0.384 | 0.701 |
| SemCueSpecific * FinalWordAlpha1Pow | 0.973 | 0.042 | 0.896 – 1.057 | -0.648 | 0.517 |
| ProbeRight * FinalWordAlpha1Conn | 1.031 | 0.055 | 0.926 – 1.148 | 0.556 | 0.578 |
| ProbeRight * RestAlpha1Conn | 0.873 | 0.071 | 0.760 – 1.004 | -1.909 | 0.056 |
| ProbeRight * FinalWordAlpha1Pow | 0.995 | 0.060 | 0.885 – 1.119 | -0.081 | 0.935 |
| FinalWordAlpha1Conn * FinalWordAlpha1Pow | 0.980 | 0.030 | 0.925 – 1.039 | -0.674 | 0.500 |
| FinalWordAlpha1Conn * RestAlpha1Conn | 0.940 | 0.031 | 0.885 – 0.998 | -2.024 | 0.043 |
| RestAlpha1Conn * FinalWordAlpha1Pow | 1.007 | 0.041 | 0.929 – 1.091 | 0.159 | 0.874 |
| Block * FinalWordAlpha1Conn | 0.912 | 0.022 | 0.874 – 0.951 | -4.294 | <0.001 |
| Block * FinalWordAlpha1Pow | 1.013 | 0.022 | 0.970 – 1.058 | 0.582 | 0.560 |
| Random Effects | | | | | |
| $\sigma^2$ | 3.2899 | | | | |
| $\tau_{00\ Pair}$ | 0.5632 | | | | |
| $\tau_{00\ Subject}$ | 0.6546 | | | | |
| $\tau_{11\ Subject.AttnCueSelective}$ | 0.2267 | | | | |
| $\tau_{11\ Subject.ProbeRight}$ | 0.3631 | | | | |
| $\rho_{01\ Subject.AttnCueSelective}$ | 0.0160 | | | | |
| $\rho_{01\ Subject.ProbeRight}$ | -0.1643 | | | | |
| ICC | 0.2932 | | | | |
| $N_{Subject}$ | 154 | | | | |
| $N_{Pair}$ | 240 | | | | |
| Observations | 35605 | | | | |
| Marginal $R^2$ / Conditional $R^2$ | 0.129 / 0.385 | | | | |

**Table S2** Summary table of the linear mixed-effects model predicting individuals' response speed. In this model brain regressors were based on the data during final word period and within $\alpha_1$ band. Significant effects after FDR-correction for multiple comparisons across model terms are shown in blue. The main effects of and interactions between listening cues are visualized in Figure 2 (main text). $\beta$: slope parameter estimate; $\sigma^2$: within-group variance; $\tau_{00}$: between-group variance; $\rho_{01}$: random-slope-intercept-correlation.

| | Speed [z] | | | | |
|---|---|---|---|---|---|
| Predictors | Estimates | std. Error | CI | Statistic | $p_{FDR}$ |
| (Intercept) | -0.034 | 0.039 | -0.111 – 0.043 | -0.869 | 0.481 |
| AttnCueSelective | 0.573 | 0.039 | 0.497 – 0.649 | 14.782 | <0.001 |
| SemCueSpecific | 0.199 | 0.032 | 0.137 – 0.261 | 6.283 | <0.001 |
| ProbeRight | 0.086 | 0.014 | 0.060 – 0.113 | 6.385 | <0.001 |
| Age | -0.154 | 0.031 | -0.215 – -0.093 | -4.948 | <0.001 |
| PTA | -0.050 | 0.031 | -0.111 – 0.011 | -1.602 | 0.273 |
| Block | 0.112 | 0.004 | 0.103 – 0.121 | 25.186 | <0.001 |
| FinalWordAlpha1Conn | -0.015 | 0.007 | -0.029 – 0.000 | -1.951 | 0.159 |
| RestAlpha1Conn | -0.052 | 0.036 | -0.123 – 0.019 | -1.432 | 0.292 |
| FinalWordAlpha1Pow | 0.014 | 0.011 | -0.008 – 0.037 | 1.264 | 0.368 |
| AttnCueSelective : SemCueSpecific | 0.091 | 0.063 | -0.033 – 0.215 | 1.435 | 0.292 |
| AttnCueSelective * FinalWordAlpha1Conn | -0.020 | 0.013 | -0.044 – 0.005 | -1.555 | 0.273 |
| AttnCueSelective * RestAlpha1Conn | -0.010 | 0.025 | -0.059 – 0.038 | -0.406 | 0.804 |
| AttnCueSelective * FinalWordAlpha1Pow | 0.018 | 0.015 | -0.012 – 0.048 | 1.190 | 0.390 |
| SemCueSpecific * FinalWordAlpha1Conn | -0.019 | 0.009 | -0.037 – -0.0004 | -2.004 | 0.159 |
| SemCueSpecific * RestAlpha1Conn | 0.003 | 0.010 | -0.016 – 0.022 | 0.284 | 0.804 |
| SemCueSpecific * FinalWordAlpha1Pow | 0.002 | 0.009 | -0.015 – 0.020 | 0.248 | 0.804 |
| ProbeRight * FinalWordAlpha1Conn | 0.003 | 0.012 | -0.019 – 0.026 | 0.288 | 0.804 |
| ProbeRight * RestAlpha1Conn | 0.016 | 0.015 | -0.013 – 0.045 | 1.094 | 0.403 |
| ProbeRight * FinalWordAlpha1Pow | 0.011 | 0.013 | -0.013 – 0.036 | 0.896 | 0.481 |
| FinalWordAlpha1Conn * FinalWordAlpha1Pow | 0.002 | 0.006 | -0.011 – 0.015 | 0.307 | 0.804 |
| FinalWordAlpha1Conn * RestAlpha1Conn | -0.008 | 0.007 | -0.022 – 0.006 | -1.150 | 0.391 |
| RestAlpha1Conn * FinalWordAlpha1Pow | -0.009 | 0.010 | -0.028 – 0.011 | -0.874 | 0.481 |
| Block * FinalWordAlpha1Conn | -0.021 | 0.005 | -0.030 – -0.012 | -4.412 | <0.001 |
| Block * FinalWordAlpha1Pow | 0.007 | 0.005 | -0.002 – 0.016 | 1.599 | 0.273 |
| **Random Effects** | | | | | |
| $\sigma^2$ | 0.588 | | | | |
| $\tau_{00\ Pair}$ | 0.056 | | | | |
| $\tau_{00\ Subject}$ | 0.196 | | | | |
| $\tau_{11\ Subject.AttnCueSelective}$ | 0.077 | | | | |
| $\tau_{11\ Subject.ProbeRight}$ | 0.017 | | | | |
| $\rho_{01\ Subject.AttnCueSelective}$ | 0.714 | | | | |
| $\rho_{01\ Subject.ProbeRight}$ | 0.085 | | | | |
| ICC | 0.322 | | | | |
| $N_{Subject}$ | 154 | | | | |
| $N_{Pair}$ | 240 | | | | |
| Observations | 32308 | | | | |
| Marginal $R^2$ / Conditional $R^2$ | 0.141 / 0.418 | | | | |

**Table S3** Summary table of the generalized linear mixed-effects model predicting individuals' listening accuracy. In this model brain regressors were based on the data during final word period and within $\boldsymbol{\alpha_2}$ band. Significant effects after FDR-correction for multiple comparisons across model terms are shown in blue. *OR*: Odds ratio; $\boldsymbol{\sigma^2}$: within-group variance; $\boldsymbol{\tau_{00}}$: between-group variance; $\boldsymbol{\rho_{01}}$: random-slope-intercept-correlation.

| | Accuracy | | | | |
|---|---|---|---|---|---|
| Predictors | Odds Ratios | std. Error | CI | Statistic | $p_{FDR}$ |
| (Intercept) | 20.50 | 0.09 | 17.23 – 24.39 | 34.10 | <0.001 |
| AttnCueSelective | 3.44 | 0.12 | 2.73 – 4.33 | 10.44 | <0.001 |
| SemCueSpecific | 1.11 | 0.11 | 0.90 – 1.36 | 0.94 | 0.350 |
| ProbeRight | 1.25 | 0.07 | 1.09 – 1.42 | 3.22 | 0.001 |
| Block | 1.29 | 0.02 | 1.24 – 1.34 | 12.12 | <0.001 |
| Age | 0.80 | 0.08 | 0.68 – 0.94 | -2.75 | 0.006 |
| PTA | 0.75 | 0.08 | 0.64 – 0.88 | -3.52 | <0.001 |
| FinalWordAlpha2Conn | 0.96 | 0.03 | 0.90 – 1.03 | -1.20 | 0.230 |
| RestAlpha2Conn | 0.97 | 0.07 | 0.84 – 1.12 | -0.42 | 0.675 |
| FinalWordAlpha2Pow | 0.98 | 0.05 | 0.89 – 1.08 | -0.41 | 0.679 |
| AttnCueSelective : SemCueSpecific | 1.36 | 0.21 | 0.89 – 2.07 | 1.43 | 0.154 |
| AttnCueSelective * FinalWordAlpha2Conn | 0.97 | 0.06 | 0.87 – 1.08 | -0.49 | 0.623 |
| AttnCueSelective * RestAlpha2Conn | 1.01 | 0.07 | 0.89 – 1.15 | 0.20 | 0.841 |
| AttnCueSelective * FinalWordAlpha2Pow | 0.99 | 0.06 | 0.89 – 1.11 | -0.10 | 0.924 |
| SemCueSpecific * FinalWordAlpha2Conn | 1.04 | 0.04 | 0.96 – 1.14 | 0.95 | 0.343 |
| SemCueSpecific * RestAlpha2Conn | 1.00 | 0.04 | 0.92 – 1.10 | 0.10 | 0.917 |
| SemCueSpecific * FinalWordAlpha2Pow | 0.96 | 0.04 | 0.89 – 1.04 | -0.95 | 0.343 |
| ProbeRight * FinalWordAlpha2Conn | 0.96 | 0.06 | 0.86 – 1.07 | -0.67 | 0.504 |
| ProbeRight * RestAlpha2Conn | 0.95 | 0.07 | 0.82 – 1.09 | -0.79 | 0.428 |
| ProbeRight * FinalWordAlpha2Pow | 1.02 | 0.06 | 0.91 – 1.14 | 0.31 | 0.757 |
| FinalWordAlpha2Conn * FinalWordAlpha2Pow | 1.00 | 0.03 | 0.95 – 1.07 | 0.17 | 0.868 |
| FinalWordAlpha2Conn * RestAlpha2Conn | 0.95 | 0.03 | 0.89 – 1.02 | -1.42 | 0.156 |
| RestAlpha2Conn * FinalWordAlpha2Pow | 1.00 | 0.04 | 0.92 – 1.08 | -0.10 | 0.918 |
| Block * FinalWordAlpha2Conn | 0.92 | 0.02 | 0.88 – 0.96 | -3.74 | <0.001 |
| Block * FinalWordAlpha2Pow | 1.00 | 0.02 | 0.95 – 1.04 | -0.12 | 0.907 |
| Random Effects | | | | | |
| $\sigma^2$ | 3.29 | | | | |
| $\tau_{00\ Pair}$ | 0.56 | | | | |
| $\tau_{00\ Subject}$ | 0.66 | | | | |
| $\tau_{11\ Subject.AttnCueSelective}$ | 0.22 | | | | |
| $\tau_{11\ Subject.ProbeRight}$ | 0.37 | | | | |
| $\rho_{01\ Subject.AttnCueSelective}$ | 0.02 | | | | |
| $\rho_{01\ Subject.ProbeRight}$ | -0.14 | | | | |
| ICC | 0.29 | | | | |
| $N_{Subject}$ | 154 | | | | |
| $N_{Pair}$ | 240 | | | | |
| Observations | 35605 | | | | |
| Marginal $R^2$ / Conditional $R^2$ | 0.126 / 0.383 | | | | |

**Table S4** Summary table of the linear mixed-effects model predicting individuals' response speed. In this model brain regressors were based on the data during final word period and within $\alpha_2$ band. Significant effects after FDR-correction for multiple comparisons across model terms are shown in blue. $\beta$: slope parameter estimate; $\sigma^2$: within-group variance; $\tau_{00}$: between-group variance; $\rho_{01}$: random-slope-intercept-correlation.

| Predictors | Speed [z] Estimates | std. Error | CI | Statistic | $p_{FDR}$ |
|---|---|---|---|---|---|
| (Intercept) | -0.045 | 0.040 | -0.123 – 0.033 | -1.127 | 0.464 |
| AttnCueSelective | 0.573 | 0.039 | 0.497 – 0.649 | 14.763 | <0.001 |
| SemCueSpecific | 0.199 | 0.032 | 0.137 – 0.261 | 6.286 | <0.001 |
| ProbeRight | 0.087 | 0.014 | 0.060 – 0.113 | 6.409 | <0.001 |
| Age | -0.159 | 0.032 | -0.221 – -0.097 | -5.036 | <0.001 |
| PTA | -0.053 | 0.031 | -0.114 – 0.009 | -1.681 | 0.232 |
| Block | 0.110 | 0.004 | 0.101 – 0.119 | 24.591 | <0.001 |
| FinalWordAlpha2Conn | -0.005 | 0.007 | -0.019 – 0.009 | -0.698 | 0.713 |
| RestAlpha2Conn | -0.014 | 0.037 | -0.086 – 0.059 | -0.366 | 0.893 |
| FinalWordAlpha2Pow | 0.008 | 0.011 | -0.014 – 0.030 | 0.732 | 0.713 |
| AttnCueSelective : SemCueSpecific | 0.091 | 0.063 | -0.033 – 0.215 | 1.433 | 0.321 |
| AttnCueSelective * FinalWordAlpha2Conn | -0.0002 | 0.013 | -0.025 – 0.024 | -0.016 | 0.987 |
| AttnCueSelective * RestAlpha2Conn | 0.007 | 0.025 | -0.042 – 0.055 | 0.275 | 0.928 |
| AttnCueSelective * FinalWordAlpha2Pow | 0.014 | 0.015 | -0.015 – 0.043 | 0.930 | 0.587 |
| SemCueSpecific * FinalWordAlpha2Conn | -0.024 | 0.009 | -0.042 – -0.005 | -2.484 | 0.071 |
| SemCueSpecific * RestAlpha2Conn | 0.014 | 0.009 | -0.005 – 0.032 | 1.426 | 0.321 |
| SemCueSpecific * FinalWordAlpha2Pow | 0.004 | 0.009 | -0.013 – 0.021 | 0.457 | 0.852 |
| ProbeRight * FinalWordAlpha2Conn | -0.002 | 0.012 | -0.025 – 0.021 | -0.164 | 0.942 |
| ProbeRight * RestAlpha2Conn | 0.009 | 0.014 | -0.020 – 0.037 | 0.602 | 0.760 |
| ProbeRight * FinalWordAlpha2Pow | -0.003 | 0.012 | -0.027 – 0.021 | -0.232 | 0.928 |
| FinalWordAlpha2Conn * FinalWordAlpha2Pow | -0.008 | 0.007 | -0.021 – 0.004 | -1.277 | 0.388 |
| FinalWordAlpha2Conn * RestAlpha2Conn | 0.012 | 0.007 | -0.002 – 0.027 | 1.716 | 0.232 |
| RestAlpha2Conn * FinalWordAlpha2Pow | -0.030 | 0.010 | -0.049 – -0.011 | -3.122 | 0.060 |
| Block * FinalWordAlpha2Conn | -0.032 | 0.005 | -0.041 – -0.023 | -6.693 | <0.001 |
| Block * FinalWordAlpha2Pow | 0.001 | 0.005 | -0.009 – 0.010 | 0.120 | 0.942 |
| **Random Effects** | | | | | |
| $\sigma^2$ | 0.588 | | | | |
| $\tau_{00\ Pair}$ | 0.056 | | | | |
| $\tau_{00\ Subject}$ | 0.205 | | | | |
| $\tau_{11\ Subject.AttnCueSelective}$ | 0.077 | | | | |
| $\tau_{11\ Subject.ProbeRight}$ | 0.017 | | | | |
| $\rho_{01\ Subject.AttnCueSelective}$ | 0.725 | | | | |
| $\rho_{01\ Subject.ProbeRight}$ | 0.079 | | | | |
| ICC | 0.329 | | | | |
| $N_{Subject}$ | 154 | | | | |
| $N_{Pair}$ | 240 | | | | |
| Observations | 32308 | | | | |
| Marginal $R^2$ / Conditional $R^2$ | 0.138 / 0.422 | | | | |

**Table S5** Summary table of the generalized linear mixed-effects model predicting individuals' listening accuracy. In this model brain regressors were based on the data during spatial cue and within $\boldsymbol{\beta_1}$ band. Significant effects after FDR-correction for multiple comparisons across model terms are shown in blue. *OR*: Odds ratio; $\boldsymbol{\sigma^2}$: within-group variance; $\boldsymbol{\tau_{00}}$: between-group variance; $\boldsymbol{\rho_{01}}$: random-slope-intercept-correlation.

| | Accuracy | | | | |
|---|---|---|---|---|---|
| Predictors | Odds Ratios | std. Error | CI | Statistic | $p_{FDR}$ |
| (Intercept) | 20.07 | 0.09 | 16.93 – 23.80 | 34.57 | <0.001 |
| AttnCueSelective | 3.44 | 0.12 | 2.73 – 4.33 | 10.45 | <0.001 |
| SemCueSpecific | 1.11 | 0.11 | 0.90 – 1.36 | 0.93 | 0.351 |
| ProbeRight | 1.24 | 0.07 | 1.09 – 1.42 | 3.17 | 0.002 |
| Block | 1.27 | 0.02 | 1.22 – 1.32 | 11.81 | <0.001 |
| Age | 0.81 | 0.08 | 0.69 – 0.94 | -2.66 | 0.008 |
| PTA | 0.75 | 0.08 | 0.64 – 0.87 | -3.69 | <0.001 |
| AttnCueBeta1Conn | 1.04 | 0.03 | 0.98 – 1.10 | 1.41 | 0.159 |
| RestBeta1Conn | 0.94 | 0.07 | 0.82 – 1.07 | -0.94 | 0.350 |
| AttnCueBeta1Pow | 0.96 | 0.04 | 0.89 – 1.04 | -1.00 | 0.316 |
| AttnCueSelective : SemCueSpecific | 1.36 | 0.21 | 0.89 – 2.07 | 1.43 | 0.153 |
| AttnCueSelective * AttnCueBeta1Conn | 0.97 | 0.05 | 0.88 – 1.07 | -0.67 | 0.503 |
| AttnCueSelective * RestLowBetaConn | 0.99 | 0.06 | 0.88 – 1.11 | -0.19 | 0.851 |
| AttnCueSelective * AttnCueBeta1Pow | 0.96 | 0.06 | 0.86 – 1.07 | -0.66 | 0.507 |
| ProbeRight * AttnCueBeta1Conn | 0.92 | 0.05 | 0.84 – 1.01 | -1.71 | 0.088 |
| ProbeRight * RestBeta1Conn | 0.97 | 0.07 | 0.85 – 1.11 | -0.44 | 0.658 |
| ProbeRight * AttnCueBeta1Pow | 1.04 | 0.06 | 0.93 – 1.16 | 0.64 | 0.523 |
| AttnCueBeta1Conn * AttnCueBeta1Pow | 0.99 | 0.03 | 0.94 – 1.04 | -0.42 | 0.673 |
| AttnCueBeta1Conn * RestBeta1Conn | 0.95 | 0.03 | 0.90 – 1.01 | -1.77 | 0.076 |
| RestBeta1Conn * AttnCueBeta1Pow | 0.99 | 0.04 | 0.92 – 1.06 | -0.33 | 0.744 |
| Block * AttnCueBeta1Conn | 0.95 | 0.02 | 0.91 – 0.99 | -2.32 | 0.020 |
| Block * AttnCueBeta1Pow | 0.97 | 0.02 | 0.92 – 1.01 | -1.47 | 0.140 |
| Random Effects | | | | | |
| $\sigma^2$ | 3.29 | | | | |
| $\tau_{00\ Pair}$ | 0.56 | | | | |
| $\tau_{00\ Subject}$ | 0.65 | | | | |
| $\tau_{11\ Subject.AttnCueSelective}$ | 0.22 | | | | |
| $\tau_{11\ Subject.ProbeRight}$ | 0.37 | | | | |
| $\rho_{01\ Subject.AttnCueSelective}$ | 0.03 | | | | |
| $\rho_{01\ Subject.ProbeRight}$ | -0.13 | | | | |
| ICC | 0.29 | | | | |
| $N_{Subject}$ | 154 | | | | |
| $N_{Pair}$ | 240 | | | | |
| Observations | 35605 | | | | |
| Marginal $R^2$ / Conditional $R^2$ | 0.126 / 0.381 | | | | |

**Table S6** Summary table of the linear mixed-effects model predicting individuals' response speed. In this model brain regressors were based on the data during spatial cue period and within $\boldsymbol{\beta_1}$ band. Significant effects after FDR-correction for multiple comparisons across model terms are shown in blue. The main effects of and interactions between listening cues are visualized in Figure 2 (main text). The interaction between rest and task connectivity is visualized in Figure 5 (main text). β: slope parameter estimate; $\boldsymbol{\sigma^2}$: within-group variance; $\boldsymbol{\tau_{00}}$: between-group variance; $\boldsymbol{\rho_{01}}$: random-slope-intercept-correlation.

| | Speed [z] | | | | |
|---|---|---|---|---|---|
| Predictors | Estimates | std. Error | CI | Statistic | $p_{FDR}$ |
| (Intercept) | -0.042 | 0.039 | -0.119 – 0.035 | -1.064 | 0.372 |
| AttnCueSelective | 0.573 | 0.039 | 0.497 – 0.649 | 14.747 | <0.001 |
| SemCueSpecific | 0.199 | 0.032 | 0.137 – 0.261 | 6.276 | <0.001 |
| ProbeRight | 0.086 | 0.013 | 0.060 – 0.112 | 6.443 | <0.001 |
| Age | -0.147 | 0.031 | -0.208 – -0.086 | -4.702 | <0.001 |
| PTA | -0.060 | 0.031 | -0.121 – 0.001 | -1.925 | 0.149 |
| Block | 0.108 | 0.004 | 0.100 – 0.117 | 24.939 | <0.001 |
| AttnCueBeta1Conn | -0.010 | 0.006 | -0.022 – 0.002 | -1.610 | 0.185 |
| RestBeta1Conn | -0.017 | 0.037 | -0.089 – 0.054 | -0.474 | 0.736 |
| AttnCueBeta1Pow | -0.029 | 0.009 | -0.046 – -0.012 | -3.276 | 0.004 |
| AttnCueSelective : SemCueSpecific | 0.091 | 0.063 | -0.033 – 0.216 | 1.444 | 0.218 |
| AttnCueSelective * AttnCueBeta1Conn | -0.001 | 0.011 | -0.023 – 0.020 | -0.134 | 0.893 |
| AttnCueSelective * RestBeta1Conn | 0.005 | 0.024 | -0.043 – 0.052 | 0.188 | 0.893 |
| AttnCueSelective * AttnCueBeta1Pow | -0.022 | 0.014 | -0.048 – 0.005 | -1.583 | 0.185 |
| ProbeRight * AttnCueBeta1Conn | -0.019 | 0.010 | -0.039 – 0.001 | -1.817 | 0.169 |
| ProbeRight * RestBeta1Conn | 0.016 | 0.014 | -0.011 – 0.043 | 1.181 | 0.327 |
| ProbeRight * AttnCueBeta1Pow | -0.002 | 0.012 | -0.024 – 0.021 | -0.136 | 0.893 |
| AttnCueBeta1Conn * AttnCueBeta1Pow | 0.004 | 0.006 | -0.007 – 0.015 | 0.683 | 0.604 |
| AttnCueBeta1Conn * RestBeta1Conn | 0.019 | 0.006 | 0.007 – 0.031 | 3.108 | 0.006 |
| RestBeta1Conn * AttnCueBeta1Pow | 0.015 | 0.009 | -0.002 – 0.032 | 1.757 | 0.174 |
| Block * AttnCueBeta1Conn | -0.007 | 0.005 | -0.016 – 0.002 | -1.600 | 0.185 |
| Block * AttnCueBeta1Pow | 0.007 | 0.005 | -0.002 – 0.016 | 1.566 | 0.185 |
| Random Effects | | | | | |
| $\sigma^2$ | 0.588 | | | | |
| $\tau_{00\ Pair}$ | 0.056 | | | | |
| $\tau_{00\ Subject}$ | 0.201 | | | | |
| $\tau_{11\ Subject.AttnCueSelective}$ | 0.078 | | | | |
| $\tau_{11\ Subject.ProbeRight}$ | 0.016 | | | | |
| $\rho_{01\ Subject.AttnCueSelective}$ | 0.719 | | | | |
| $\rho_{01\ Subject.ProbeRight}$ | 0.072 | | | | |
| ICC | 0.326 | | | | |
| $N_{Subject}$ | 154 | | | | |
| $N_{Pair}$ | 240 | | | | |
| Observations | 32308 | | | | |
| Marginal $R^2$ / Conditional $R^2$ | 0.136 / 0.418 | | | | |
